# PNPLA3(148M) promotes hepatic steatosis by interfering with triglyceride hydrolysis through a gain-of-function mechanism

**DOI:** 10.1101/2024.08.02.606015

**Authors:** Yang Wang, Sen Hong, Hannah Hudson, Nora Kory, Lisa N. Kinch, Julia Kozlitina, Jonathan C. Cohen, Helen H. Hobbs

## Abstract

**Background & Aims:** PNPLA3(148M) (patatin-like phospholipase domain-containing protein 3) is the most impactful genetic risk factor for steatotic liver disease (SLD), thus motivating a search for therapeutic modulators of its expression. A key unresolved issue is whether PNPLA3(148M) confers a loss- or gain-of-function. Here we used multiple approaches to further test the hypothesis that PNPLA3 causes steatosis by sequestering ABHD5 (α/β hydrolase domain containing protein 5), the cofactor of ATGL (adipose TG lipase), thus limiting mobilization of hepatic triglyceride (TG).

**Methods:** We quantified the physical interactions between ABHD5 and PNPLA3/ATGL in cultured hepatocytes using NanoBiT complementation assays. Immunocytochemistry was used to compare the relative binding of PNPLA3 and ATGL to ABHD5 and to determine if PNPLA3 must associate with lipid droplets (LDs) to inhibit ATGL. Adenoviruses and adeno-associated viruses were used to express PNPLA3 in liver-specific *Atgl^-/-^* mice and ABHD5 in livers of *Pnpla3^148M/M^* mice, respectively. We used purified recombinant proteins to compare the TG hydrolytic activities of PNPLA3 and ATGL in the presence and absence of ABHD5.

**Results:** ABHD5 interacted preferentially with PNPLA3 relative to ATGL in cultured hepatocytes and *in vitro,* with no differences observed between PNPLA3(WT) or PNPLA3(148M). PNPLA3(148M)-associated inhibition of TG hydrolysis required localization of PNPLA3 to LDs and the presence of ATGL. Finally, overexpression of ABHD5 reversed the hepatic steatosis in *Pnpla3^M/M^* mice.

**Conclusions:** These findings support the premise that PNPLA3(148M) promotes hepatic steatosis by accumulating on LDs and inhibiting ATGL-mediated lipolysis in an ABHD5-dependent manner. Our results predict that reducing, rather that increasing PNPLA3 expression will be the best strategy to treat PNPLA3(148M)-associated SLD.

**Impact and implications:** Steatotic liver disease (SLD) is a common complex disorder associated with both environmental and genetic risk factors. PNPLA3(148M) is the most impactful genetic risk factor for SLD and yet its pathogenic mechanism remains controversial. Here we provide evidence that PNPLA3(148M) promotes triglyceride (TG) accumulation by sequestering ABHD5, thus limiting its availability to activate ATGL. Although the substitution of methionine for isoleucine reduces the TG hydrolytic activity of PNPLA3, the loss-of-function is only indirectly related to the steatotic effect of the variant. Here we provide evidence that PNPLA3(148M) confers a gain-of-function by interfering with ATGL-mediated TG hydrolysis. These findings have implications for the design of potential PNPLA3-based therapies. Reducing, rather than increasing, PNPLA3 levels is predicted to reverse steatosis in susceptible individuals.

**Graphical Abstract:** 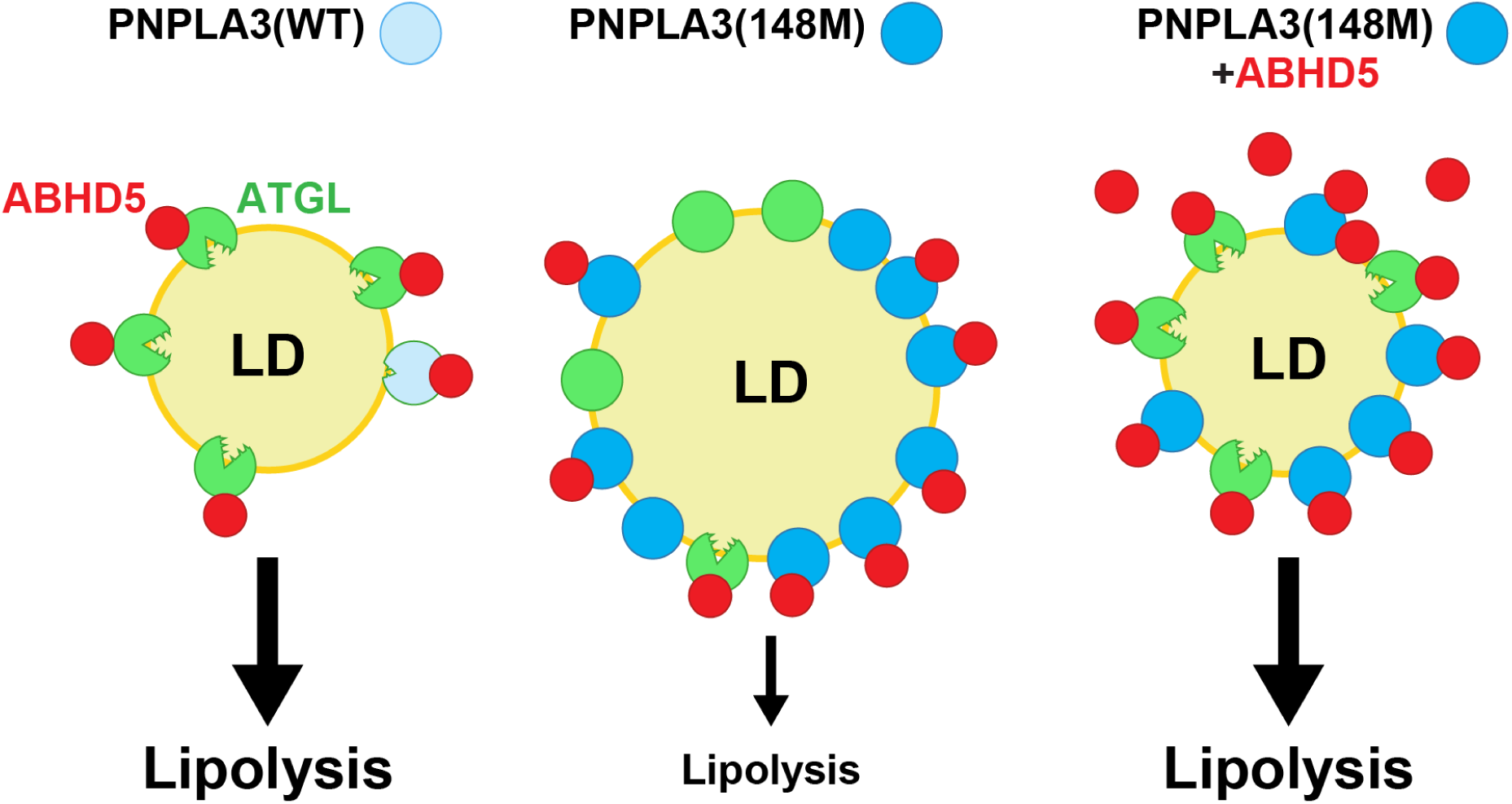

**Highlights:**

- ABHD5 binds preferentially to PNPLA3 relative to ATGL.
- PNPLA3(WT) and PNPLA3(148M) compete similarly for binding and inhibition of ATGL.
- ABHD5 activates the triglyceride lipase activity of PNPLA3, as well as ATGL.
- The steatotic effect of PNPLA3(148M) requires expression of ATGL.
- Overexpression of ABHD5 can rescue the steatosis associated with PNPLA3(148M).

## Introduction

Steatotic liver disease (SLD) is an exemplar of a complex disease. The cardinal risk factor for SLD is obesity, but nonlinear interactions among other genetic and nongenetic factors, most notably insulin resistance and excessive alcohol consumption, contribute significantly to the development and progression of this disorder.^1^ The most impactful DNA sequence variant conferring risk of SLD is a missense variant (I148M) in PNPLA3.^2, 3^ The substitution of methionine for isoleucine at codon 148 of PNPLA3 is associated with elevated hepatic TG content and progression of steatosis to steatohepatitis, cirrhosis, and hepatocellular carcinoma.^1, 2^ Despite its importance as a risk factor, the pathogenic mechanism responsible for association of PNPLA3(148M) with liver disease remains unclear, thus limiting the development of therapeutic interventions.

PNPLA3 has TG hydrolase activity,^4^ which is reduced ∼80% by the I148M substitution.^5^ Based on this finding, we initially hypothesized that PNPLA3(148M) confers a loss-of-function, causing steatosis due to the lack of TG hydrolase activity. This premise was recently also suggested by others,^6^ but is contradictory to the finding that mice lacking PNPLA3 fail to develop hepatic steatosis, even when challenged with a high-sucrose diet to promote TG synthesis.^7, 8^ Moreover, PNPLA3 does not cause steatosis unless the I148M variant is expressed, which is more consistent with the protein conferring a gain of function.^9^ Treatments that reduce PNPLA3 expression such as anti-PNPLA3 siRNAs, reverse the hepatic steatosis in mice expressing PNPLA3(148M).^10, 11^ Moreover, a recent Phase 1 study in humans reported up to a 46% reduction of hepatic TG content in PNPLA3(148M) homozygotes treated with an siRNA targeting hepatic PNPLA3.^12^

How does PNPLA3(148M) confer a gain-of-function? Previously, we showed that overexpression of PNPLA3(WT or 148M) in cultured hepatocytes inhibits ATGL-mediated TG hydrolysis.^13^ PNPLA3 most closely resembles PNPLA2, also called ATGL, the major TG lipase in most tissues.^14, 15^ Humans (and mice) lacking ATGL have ectopic accumulation of TG in multiple tissues, including the liver.^15–17^ Ectopic accumulation of TG is also a prominent feature in mice and humans lacking the co-factor of ATGL, ABHD5, also known as CGI-58 (comparative gene identification-58).^18, 19^ Addition of ABHD5 to ATGL increases its lipolytic activity 4-20 fold.^18^ The mechanism by which ABHD5 promotes ATGL activity remains unclear. Nonetheless, liver-specific inactivation of ABHD5 in mice is sufficient to cause hepatic steatosis with accompanying inflammation and fibrosis.^20^

We have speculated that PNPLA3 may limit ATGL activity by sequestering ABHD5.^13^ In support of this model, we found that the steatotic effect of PNPLA3(148M) requires expression of ABHD5.^13^ Here we provide additional compelling evidence that PNPLA3(148M) accumulation on LDs causes hepatic steatosis by limiting the access of ABHD5 to ATGL.^13^ Based on our findings, we predict that efforts directed at lowering PNPLA3 expression, either through weight reduction or by using other therapeutics, would be the most effective approach to treat PNPLA3(148M)-associated steatotic liver disease.

## Material and methods

The materials and methods used in this paper are detailed in the *Supplementary Data*.

## Results

### ABHD5 preferentially binds PNPLA3 relative to ATGL in cultured hepatocytes

Previously, we showed that purified PNPLA3 physically interacts with ABHD5, albeit weakly.^13^ Here we used the NanoBiT complementation assay system to assess more quantitatively the relative strength of the interactions between ABHD5 and both PNPLA3 and ATGL.^21^ In this assay, the nanoluciferase enzyme is divided into two parts: Large-BiT (LgBiT, L), which comprises the bulk of the enzyme (17.6 kDa) and Small-BiT (SmBiT, S), an 11-residue peptide (Fig. 1A) that complements Lg-BiT to restore luciferase activity. The two luciferase subunits have low mutual affinity, so nanoluciferase activity is reconstituted only if the proteins linked to the two subunits physically interact. We fused SmBiT to the N-terminus of PNPLA3-V5 or ATGL-V5 (S-PNPLA3, S-ATGL) and LgBiT to the C-terminus of ABHD5 (ABHD5-L) (Fig. 1A). Luciferase activity was monitored by adding a cell permeable substrate, furimazine, to the transfected cells and measuring conversion to luminescent furimamide.

**Fig. 1.**
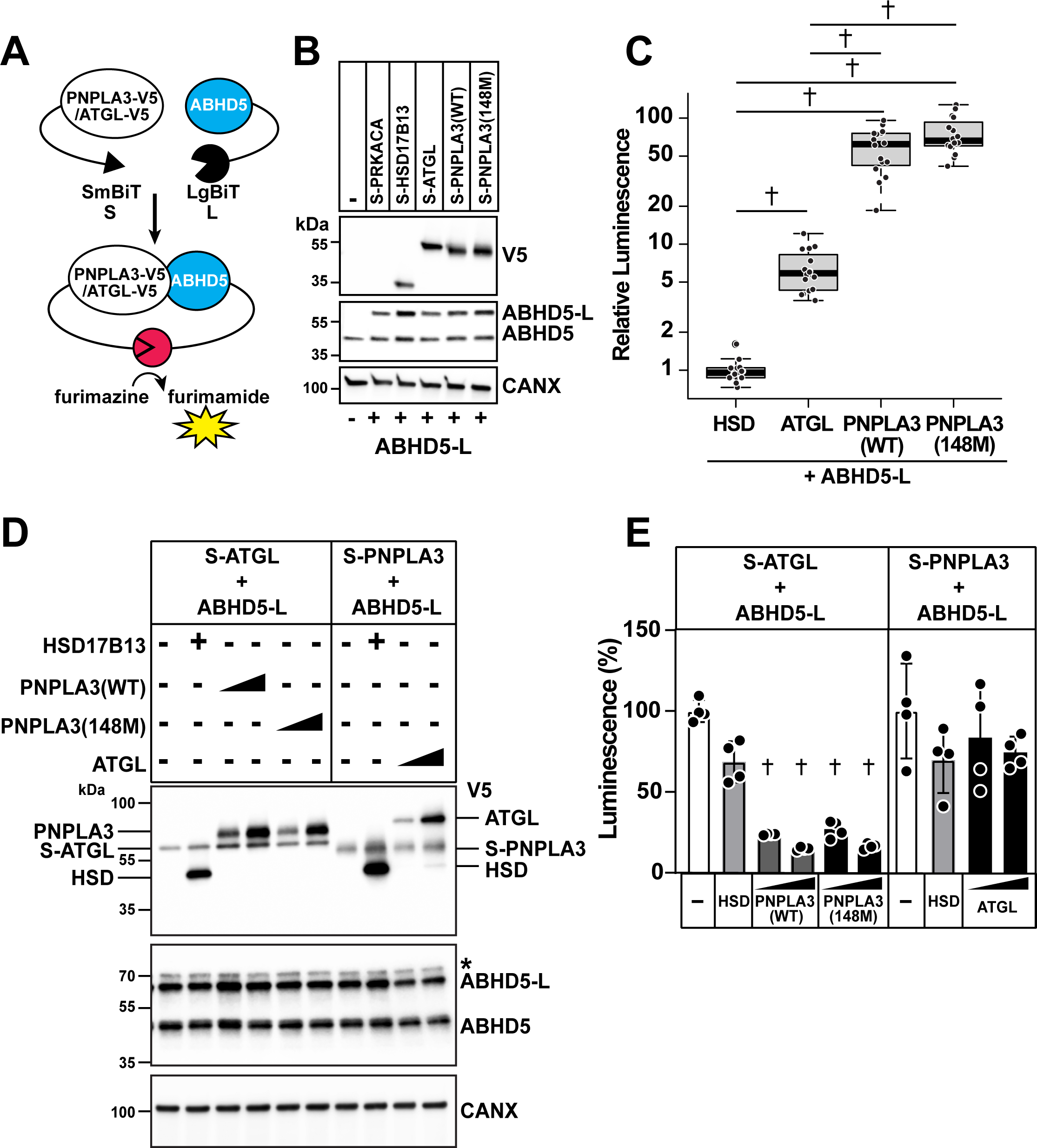
Relative strength of interactions between ABHD5 and ATGL, PNPLA3(WT or 148M) in HuH7 cells. (A) Schematic of NanoBiT protein-protein interaction assay. Small-BiT (SmBiT, S) was fused to the N-terminus of PNPLA3-V5 (S-PNPLA3) or ATGL-V5 (S-ATGL) and Large-BiT (LgBiT, L) was fused to the C-terminus of ABHD5 (ABHD5-L). Physical proximity due to protein-protein interactions between ABHD5 and PNPLA3 or ABHD5 and ATGL results in S and L forming a functional luciferase, which catalyzes conversion of furimazine to produce a luminescent signal in live cells detected by EnVision Multilabel Plate Readers. (B) NanoBiT protein-protein interaction assays were performed in HuH7 cells expressing ABHD5-L and the indicated recombinant proteins. S-PRKACA (protein kinase cAMP-activated catalytic subunit alpha), a soluble cytoplasmic protein, was used as a negative control. Immunoblot analysis of the transfected cells is shown. (C) Cellular luminescent signals were normalized to levels of the SmBiT-linked proteins, where levels of S-HSD17B13 and ABHD5-L pair were set to 1 (right). Data represent median [25^th^% - 75^th^%] of data pooled from 3 experiments using box-and-whisker plot in a natural logarithmic scale; p values of natural logarithms of relative luminescence were determined by one-way ANOVA followed by Tukey’s multiple comparisons test; †p < 0.0001. (D) NanoBiT-tagged protein pairs were co-expressed with proteins without NanoBiT tags to assess competitive binding to ABHD5-L. The competitor proteins were linked with 3×HA, 3×FLAG and 1×V5 tags at the C-terminus. The ratios of the plasmids encoding the two NanoBiT components and competitors were 1:1:0.5 or 1:1:1. The asterisk indicates a non-specific signal. (E) Luminescence signals were quantified and normalized to levels of SmBiT-linked protein. The normalized luminescence signals were compared to indicated SmBiT-linked protein and ABHD5-L pair with no competitors (set at 100%). Data represent mean ± SD of technical replicates; p values were determined by one-way ANOVA followed by Sidak’s multiple comparisons test; †p < 0.0001. The experiments were repeated three times and the results were similar.

To minimize artifacts due to use of epitope tags, we tested positional effects of SmBiT and LgBiT on signal generation and protein localization. Optimal signals were obtained when SmBiT was placed at the N-terminus of PNPLA3 (S-PNPLA3), and LgBiT was fused to the C-terminus of ABHD5 (ABHD5-L) (Fig. S1A). Addition of SmBiT to PNPLA3 did not alter protein localization to LDs (Fig. S1B). No LDs were present in cells coexpressing S-PNPLA3(WT) and ABHD5 (Fig. S1C), whereas LDs were retained in cells expressing catalytically impaired S-PNPLA3(148M) plus ABHD5. These results are congruent with what we previously reported using untagged proteins.^13^

To examine the relative strengths of the interactions between ABHD5-L and the two lipases, the cofactor and enzymes were co-expressed in cultured human hepatocytes (HuH7 cells) and luminescence was measured after 24 h. A cytoplasmic protein (S-PRKACA, Protein Kinase cAMP-Activated Catalytic Subunit Alpha) was used as a negative control and HSD17B13 (Hydroxysteroid 17-β dehydrogenase 13), a structurally unrelated LD protein, was included in the experiment for comparison. An immunoblot of lysates from the transfected cells is provided in Fig. 1B. Protein levels were normalized and then corrected for levels of S-protein expression. The mean value obtained for ABHD5-L plus S-HSD was set to one (Fig. 1C) and the median [25^th^-75^th^ percentile] luminescence of the interaction of ABHD5-L and S-ATGL was 4-fold higher. The median luminescence seen with co-expression of S-PNPLA3(WT or 148M) and ABHD5-L was 15-fold higher than that seen between S-ATGL and its co-factor (note: logarithmic scale on y-axis). No significant difference was found between the luminescence signal generated in cells expressing ABHD5-L and either S-PNPLA3(WT) or S-PNPLA3(148M).

To further characterize the physical interactions between ABHD5 and the two structurally related lipases, we performed a competition assay. S-ATGL and ABHD5-L were co-expressed in cells alone or together with HSD17B13, PNPLA3(WT) or PNPLA3(148M). Epitope tags were added to the C-terminus so proteins migrated different distances upon electrophoresis (Fig. 1D). A nonsignificant reduction in luminescence was seen with addition of HSD17B13 (Fig. 1E). In contrast, co-expression of PNPLA3(WT or 148M) resulted in a marked (70-85%) reduction in luminescence (Fig. 1E, left). When we co-expressed S-PNPLA3 and ABHD5-L alone or together with HSD17B13 and ATGL, no significant reductions in cellular luminescence were seen (Fig. 1E, right). We concluded from these experiments that ABHD5 interacts more robustly with PNPLA3 than ATGL and that the I148M substitution did not alter the strength of the protein interaction. Therefore, the steatotic effect of PNPLA3(148M) cannot be ascribed to the risk isoform having enhanced binding to ABHD5.

### Recruitment of ABHD5 to LDs by PNPLA3 and ATGL

To further assess the relative strength of the interactions between ABHD5 and ATGL/PNPLA3, we generated two forms of ABHD5 that retain the ability to bind ATGL but do not localize to LDs, as described.^22^ The sequences required for LD binding were mapped by Oberer and her colleagues to reside in the N-terminal 30 amino acids; deletion of those residues or substitution of alanine for three tryptophans result in failure of ABHD5 to localize to LDs (Fig. 2A and 2B).^22^ We named these lipid droplet binding defective forms of ABHD5: ABHD5-LBD1 and -LBD2. We then asked if ATGL or PNPLA3 could recruit ABHD5-LBDs to LDs. Here we used *ATGL^-/-^* QBI-293A cells (Fig. S2A) and catalytically inactive lipases [PNPLA3(47A) and ATGL(47A)] so that LDs would be retained with co-expression of the lipases with ABHD5.

**Fig. 2.**
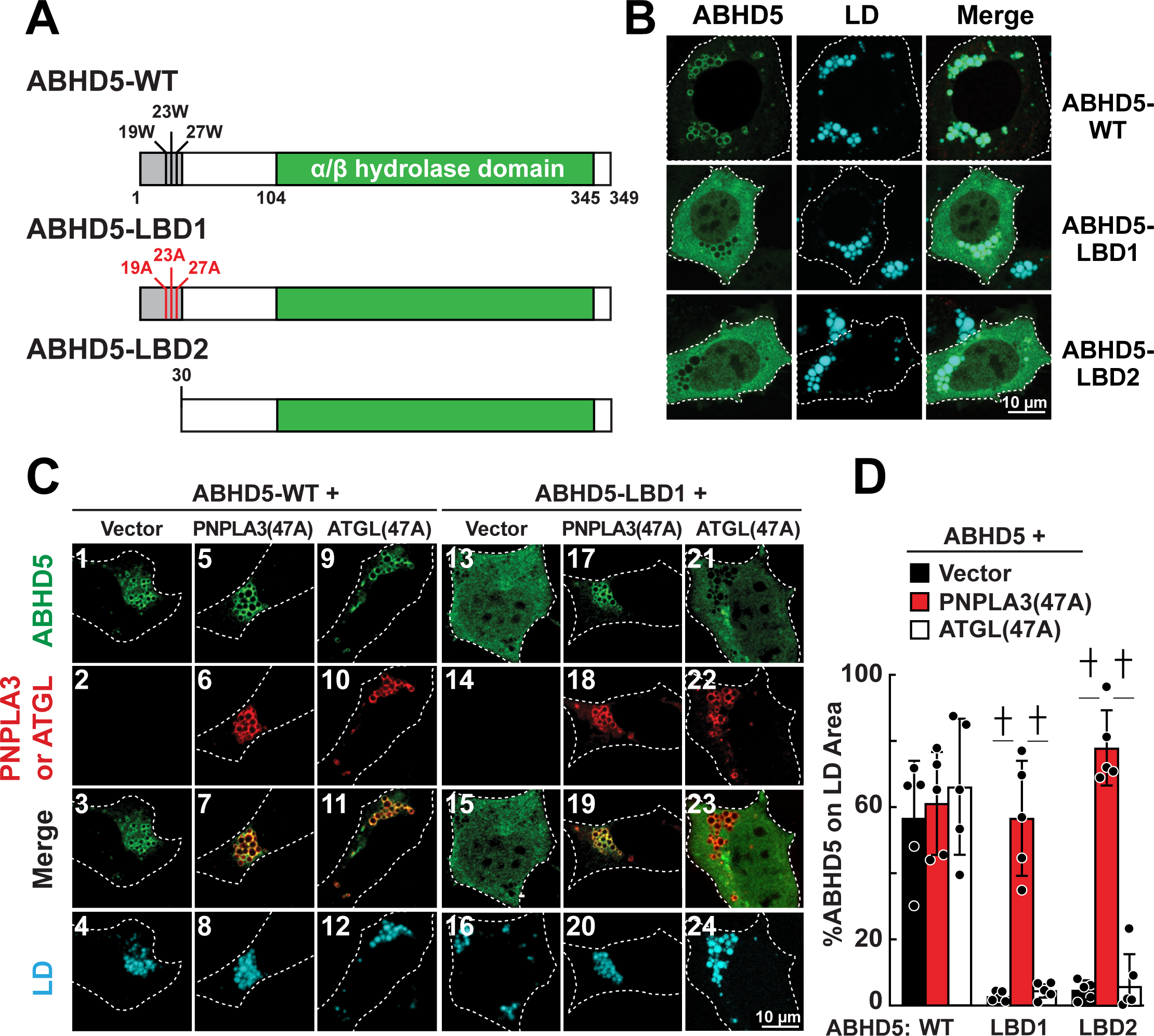
Recruitment of LD binding defective ABHD5 (ABHD5-LBD1 and ABHD5-LBD2) to LDs by PNPLA3 and ATGL in *ATGL^-/-^*QBI-293A cells. (A) Schematics of human ABHD5(WT), ABHD5(3W3A) [ABHD5-LBD1] and ABHD5(Δ30) [ABHD5-LBD2]. The first 30 residues (gray box) are required for LD binding.^22^ (B) ABHD5-WT and the LBD variants with a myc epitope tag at the C-terminus were expressed in *ATGL^-/-^* QBI-293A cells grown in medium supplemented with oleate (200 µM). Immunofluorescence was performed using a rabbit anti-myc polyclonal antibody to detect ABHD5 (green). LDs were stained using LipidTOX Deep Red (cyan). White dashes outline the transfected cells. (C) ABHD5-WT-myc or ABHD5-LBD1-myc were co-expressed with PNPLA3(47A)-V5 or ATGL(47A)-V5 in *ATGL^-/-^* cells. LDs were stained using LipidTOX Deep Red (cyan). ABHD5 (green), PNPLA3 and ATGL (red) were visualized using anti-myc and anti-V5 primary antibodies and Alexa Fluor-conjugated secondary antibodies. Merge: ABHD5 plus PNPLA3 or ATGL. Vector: pcDNA3.1(+). White dashes outline the transfected cells. (D) The proportion of ABHD5 on LDs was quantitated using ImageJ. Data represent mean ± SD; n=5 cells/group; p values were determined using 2-way ANOVA followed by Tukey’s multiple comparisons test to compare within groups; † p<0.0001. Experiments were repeated twice, and results were similar.

As expected, both inactive lipases localized to LDs, as did ABHD5 (Fig. 2C, 3, 7, 11). We then repeated the experiment using ABHD5-LBD1 and -LBD2 (Fig. 2C, left and Fig. S2B). ABHD5-LBD1 and -LBD2 were dispersed when co-expressed with empty vector (Fig. 2C, 13 and Fig. S2B, 1); less than 3% of the lipase localized to LDs (Fig. 2D). In contrast, when ABHD5-LBD was co-expressed with PNPLA3(47A), ∼60% of ABHD5-LBD1 and -LBD2 was located on LDs (Fig. 2C, 19 and Fig. S2B, 7), which was similar to the proportion seen in cells expressing wildtype (WT) co-factor (Fig. 2C, 3). A much smaller proportion of ABHD5-LBD (<6%) localized to LDs in cells expressing ATGL(47A) (Fig. 2C, 23 and Fig. S2B, 11). From these experiments, we concluded that PNPLA3 was more effective at recruiting/retaining ABHD5-LBDs to LDs than ATGL.

**Fig. 3.**
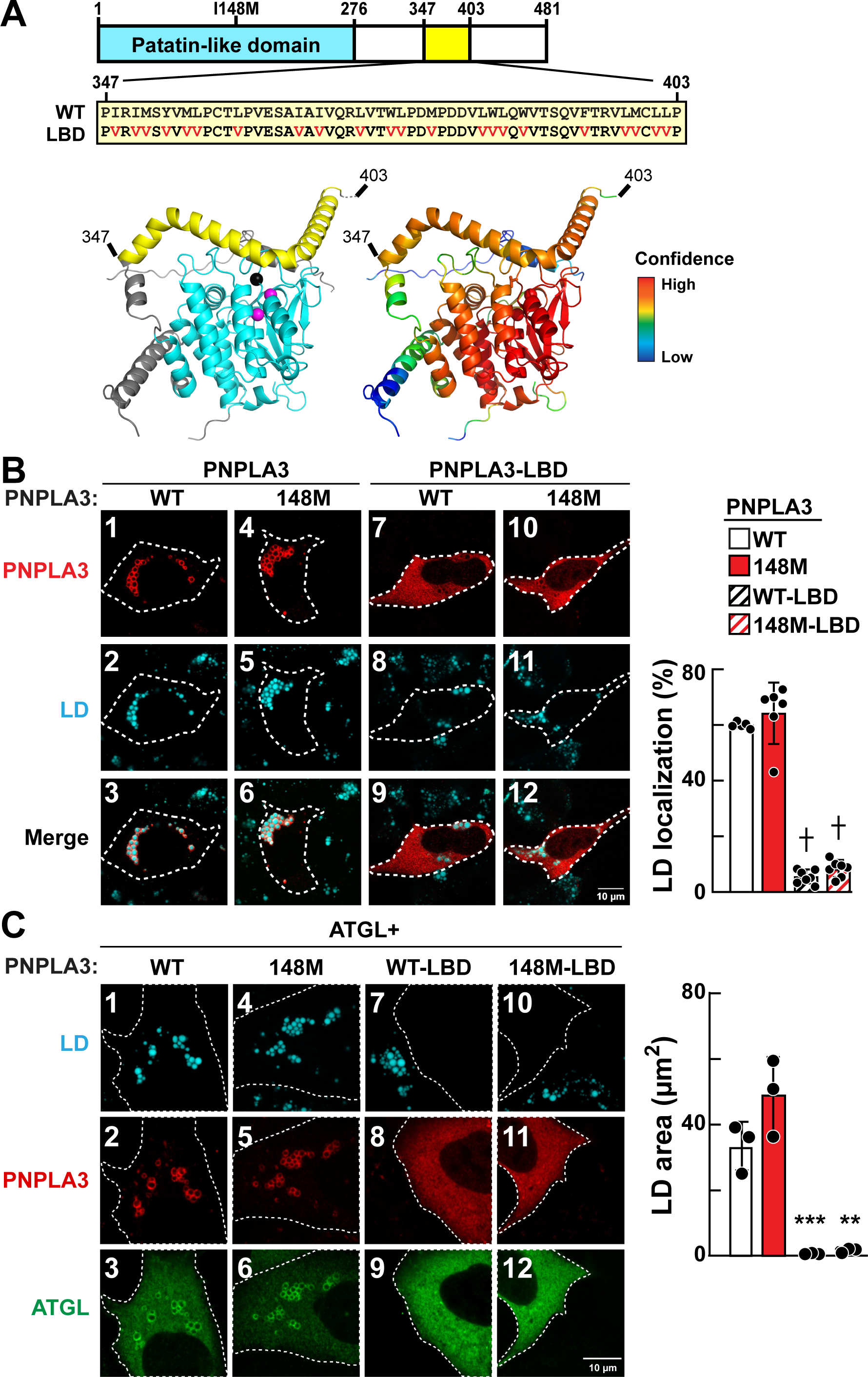
ATGL inhibition by PNPLA3 requires localization of PNPLA3 to LDs. (A) Schematic of mutations in the LD binding motif (yellow, residues 347-403) to generate the LD-binding defective (LBD) variant of PNPLA3. Valine (V) substitutions are shown in red. Human PNPLA3 structure was predicted by AlphaFold 2 (left): Yellow: putative LD binding region; Cyan: patatin-like domain; Magenta spheres: catalytic dyad (Ser47 and Asp166); Black sphere: Ile148; Grey: low confidence regions of the structure. Low confidence C-terminal region (residue 409 to 481) have been removed for clarity. (right) Scheme of human PNPLA3 colored in rainbow according to pLDDT residue pLDDT confidence: lowest pLDDT (blue) to highest pLDDT (red). (B) PNPLA3(WT) or PNPLA3(148M) +/- LBD mutations were expressed in QBI-293A cells grown in medium supplemented with oleate (200 µM). (left) Immunofluorescence was performed to show localization of PNPLA3 (red) using anti-PNPLA3 primary antibody and Alexa 555-conjugated anti-mouse secondary antibody. LDs were stained using monodansylpentane (MDH, cyan). White dashes outline transfected cells. (right) Quantification of proportion of different forms of PNPLA3 localized to LD (n=5-7 cells/group). (C) Human ATGL (green) was co-expressed with PNPLA3(WT) or PNPLA3(148M) +/- LBD mutations (red) and processed as described in Panel B, except that ATGL was visualized using an anti-ATGL primary antibody. (right) Quantification of LD area in cells co-expressing target proteins (n=3 cells/group). Data are represented as mean ± SD; p values were determined by one-way ANOVA followed by Dunnett’s multiple comparisons test; **p < 0.01, ***p< 0.001, †p<0.0001. Experiment was repeated and the results were similar.

### PNPLA3 must localize to LDs to inhibit ATGL enzymatic activity

To determine if PNPLA3 must be on LDs to inhibit ATGL activity, we developed an LBD form of PNPLA3 (PNPLA3-LBD). We identified a 57-residue region (residues 347-403) that is rich in hydrophobic amino acids and predicted to contain a long amphipathic alpha helix. The amphipathic helix rims the active site surface in the AlphaFold structure model (Fig. 3A), with the two residues comprising the catalytic site shown in magenta and residue 148 shown in black. Valine was substituted for the larger hydrophobic residues within the putative LD binding sequence. We confirmed that PNPLA3(WT and 148M)-LBD failed to localize to LDs (Fig. 3B).

Next, we tested if expression of PNPLA3-LBD inhibited ATGL-mediated TG hydrolysis. As a positive control, we co-expressed PNPLA3(WT or 148M) and ATGL in cells. Overexpression of either protein inhibited TG hydrolysis, resulting in retention of LDs (Fig. 3C, images 1 and 4).^13^ In contrast, when PNPLA3(WT or 148M)-LBD were co-expressed with ATGL, no LDs were visible in the cells (Fig. 3C, 7, 10). We concluded from these experiment that PNPLA3 must reside on LDs to inhibit the TG hydrolase activity of ATGL. We cannot rule out the possibility that PNPLA3-LBD failed to inhibit ATGL-mediated TG hydrolysis because the mutations we introduced interfered with another aspect of PNPLA3 function related to ATGL inhibition.

### PNPLA3(148M) requires expression of ATGL to promote hepatic steatosis

If PNPLA3(148M) causes steatosis by inactivating ATGL, we would expect that expression of the variant protein in *Atgl^-/-^* mice would not worsen the steatosis. We expressed both PNPLA3(WT) and PNPLA3(148M) in liver-specific *Atgl* knockout mice [(Ls)-*Atgl^-/-^* mice].^16, 23^ Immunoblot analysis of LD proteins from WT and Ls-*Atgl^-/-^* mice is shown in Fig. 4A. No differences in levels of PNPLA3 or ABHD5 were seen between the strains. We then infected the mice with adenoviruses expressing PNPLA3(WT) or PNPLA3(148M) (Fig. 4B). The ATGL levels were similar in the *Atgl^f/f^* mice receiving the control virus (RR5) or the viruses encoding the PNPLA3 isoforms (Fig. 4C). A low but detectable level of ATGL was seen in LDs from the Ls-*Atgl^-/-^* mice in this experiment, presumably due to ATGL expression in non-hepatocyte cells. Expression of PNPLA3(WT or 148M) caused a 3-fold increase in ABHD5 levels on LDs in both groups of mice without associated changes in hepatic ABHD5 mRNA levels (Fig. 4D).

**Fig. 4.**
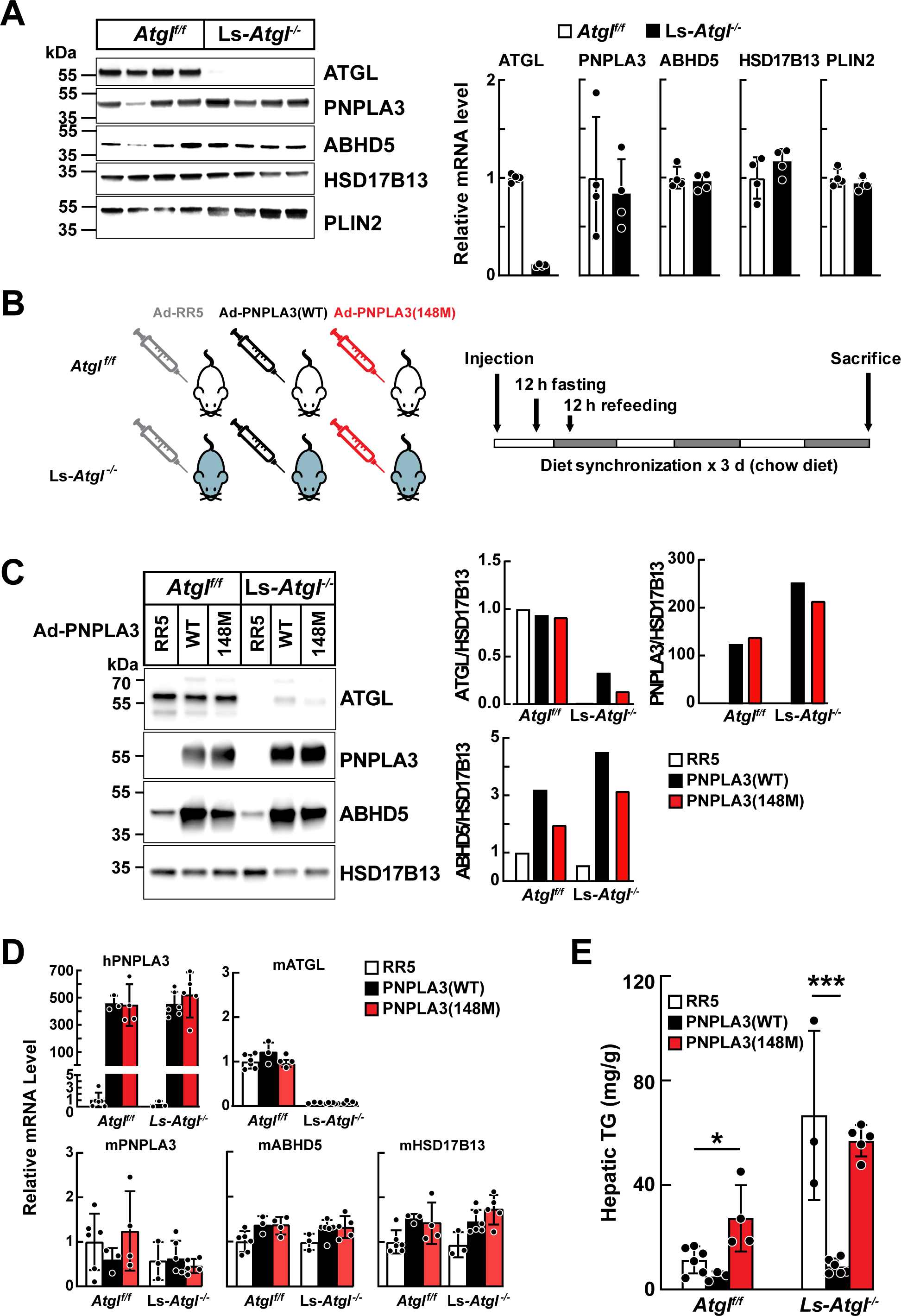
ATGL is required for PNPLA3(148M)-mediated hepatic steatosis. (A) Protein (left) and transcript levels (right) of LD proteins in Ls-*Atgl^-/-^*mice. LDs were purified from chow-fed *Atgl^f/f^* and Ls-*Atgl^-/-^*female mice (8-10 weeks, n=4 mice/group) and immunoblot analysis was performed (5 µg of protein) using the indicated antibodies (left). Total RNA was extracted from the livers and mRNA levels were measured using quantitative real-time PCR and normalized to levels of CANX mRNA. The mRNA levels in *Atgl^f/f^* mice were set as 1. The experiment was repeated with mice fed a high sucrose diet and the results were similar. (B) Schematic of experimental design. Chow-fed *Atgl^f/f^* and Ls- *Atgl^-/-^* male mice (14-17 weeks, n=3-6 mice/group) were infected with adenovirus [1.5×10^11^ genomic copies (GC)] with no insert (RR5) or expressing human PNPLA3(WT or 148M)-V5. After 3 days, diets were synchronized and mice killed at the end of the last feeding period. (C) Immunoblot analysis of pooled hepatic LD proteins (1 µg) in infected mice (left). Protein levels were quantified using LI-COR after normalization to levels of HSD17B13 (right). (D) Relative levels of hepatic mRNAs encoding LD proteins in adenovirus-infected mice. *Atgl^f/f^* mice treated with adenovirus expressing no insertion (Ad-RR5) were arbitrarily set to 1. h: human; m: mouse. (E) Hepatic TG levels were measured using an enzymatic assay and normalized to tissue weight. Data represent mean ± SD; p values were determined by one-way ANOVA followed by Tukey’s multiple comparisons test; *p<0.05, ***p<0.001. The experiment was repeated twice and the results were similar.

Hepatic TG levels were similar in groups of *Atgl^f/f^*mice infected with control virus and virus encoding PNPLA3(WT), but increased when in those expressing PNPLA3(148M) (Fig. 4E). As expected, levels of hepatic TG were higher in Ls-*Atgl^-/-^* mice and expression of PNPLA3(WT) caused a reduction in hepatic TG levels, as reported previously.^6^ Thus, PNPLA3 promotes TG lipolysis in the absence of ATGL. No further increase in hepatic TG was seen in the Ls-*Atgl^-/-^*mice that received the PNPLA3(148M)-expressing virus. Thus, expression of PNPLA3(148M) failed to promote an increase in TG accumulation in the absence of ATGL.

### Expression of ABHD5 reverses hepatic steatosis in *Pnpla3^M/M^* mice

If the amount of ABHD5 is rate-limiting in PNPLA3 knockin mice (*Pnpla3^M/M^* mice), then we would expect that the hepatic steatosis could be reversed with overexpression of ABHD5. WT and *Pnpla3^M/M^*mice were infected with AAV8 expressing ABHD5 containing three copies of a FLAG tag at the C-terminus (labeled ABHD5^#^). AAV8-GFP was used as a negative control in this experiment (Fig. 5A). As expected, hepatic levels of PNPLA3 on LDs were higher in *Pnpla3^M/M^* mice than WT mice (with no changes in PNPLA3 mRNA levels), irrespective of whether the mice received the GFP- or ABHD5-expressing virus (Fig. 5B, lanes 9-17 and Fig. 5C). Levels of PNPLA3 decreased in WT mice receiving AAV8-ABHD5^#^ (Fig. 5B, lanes 5-8 versus lanes 1-4) for reasons that are not apparent at this time.

**Fig. 5.**
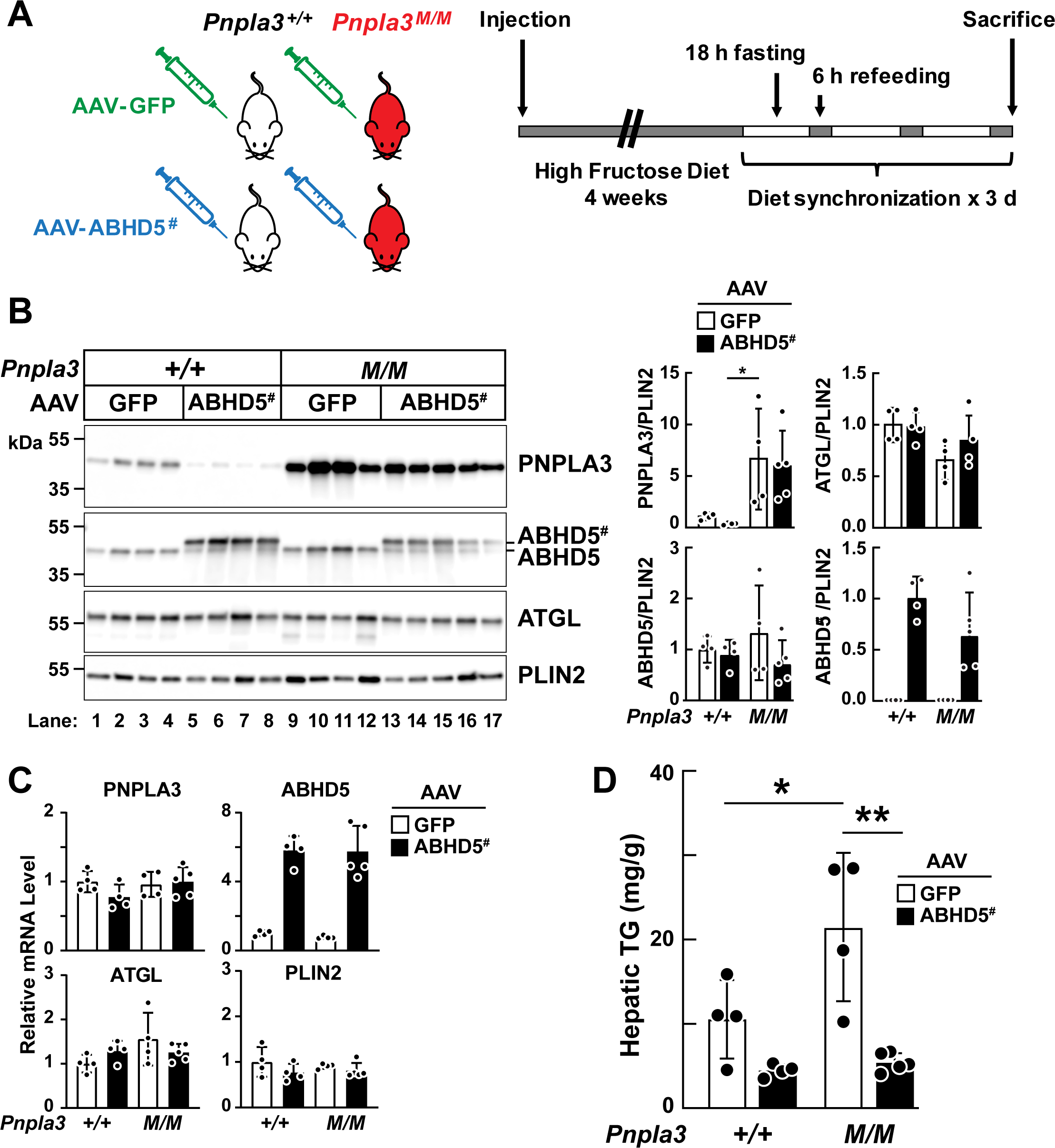
AAV-mediated hepatic expression of mouse ABHD5 rescued steatosis in *Pnpla3^M/M^* mice. (A) Schematic of experimental design. *Pnpla3^+/+^* and *Pnpla3^M/M^* female mice (12-15 weeks, n=4-5 mice/group) were infected with AAV expressing GFP or ABHD5-FLAG×3 [ABHD5^#^, 1.25×10^11^ genomic copies (GC)]. Mice were fed a high-fructose diet for 4 weeks. Diets were synchronized by 18-h fasting and 6-h refeeding for 3 days before the mice were sacrificed after last feeding cycle. (B) (left) LD proteins (1.5 µg) were subjected to immunoblotting and (right) levels of indicated proteins were quantified using LI-COR and normalized to levels of PLIN2. (C) Relative mRNA levels of indicated LD associated genes. The mRNA levels in *Pnpla3^+/+^* mice infected with AAV-GFP were set as 1. (D) Hepatic TG levels were measured as described in legend to Fig. 4. Data are represented as mean ± SD; n=4-5 mice/group; p values were determined by 2-way ANOVA followed by Tukey’s multiple comparisons test; *p<0.05, **p<0.01. The experiment was repeated and the results were similar.

As expected, hepatic TG levels were higher in *Pnpla3^M/M^*mice than WT animal that received AAV-GFP (Fig. 5C). Hepatic TG levels fell to similar low levels in the *Pnpla3^M/M^* and WT mice upon infection with AAV8-ABHD5^#^. The reduction in hepatic TG levels in *Pnpla3^M/M^*mice was not due to an increase in ATGL expression (Fig. 5B), but rather was presumed to be caused by the exogenous ABHD5^#^ activating endogenous ATGL.

### Recombinant PNPLA3 is activated by ABHD5 *in vitro*

As shown previously, co-expression of ABHD5 and PNPLA3 promotes TG hydrolysis in cultured cells.^6, 13^ This had been an unanticipated result because we had no previously found any increase of PNPLA3 enzymatic activity with addition of ABHD5 when we used lysates of cells expressing the recombinant proteins.^5^ Here we repeated the experiment but used proteins that were expressed and purified (to >95% homogeneity) from mammalian HEK293S GnTI^-^ cells (Fig. 6A).^24^ A total of 5 μg of each lipase was added to a triolein-phospholipid emulsion and after one-hour, lipids were analyzed by thin layer chromatography (TLC) (Fig. 6B). As expected, ATGL generated free fatty acids (FA), diacylglycerol (DAG) and 1-monoacylglycerol (MAG) (lane 2 vs lane 9); addition of an equimolar amount of ABHD5 caused an increase in the amounts of hydrolytic products (lane 3). A similar pattern of bands was seen with PNPLA3(WT), although the intensity of the signals was lower (lane 4). A further reduction in signal was seen when PNPLA3(148M) was used in the experiment (lane 6). Addition of ABHD5 increased the TG hydrolase activity of both forms of PNPLA3 (lanes 5 and 7).

**Fig. 6.**
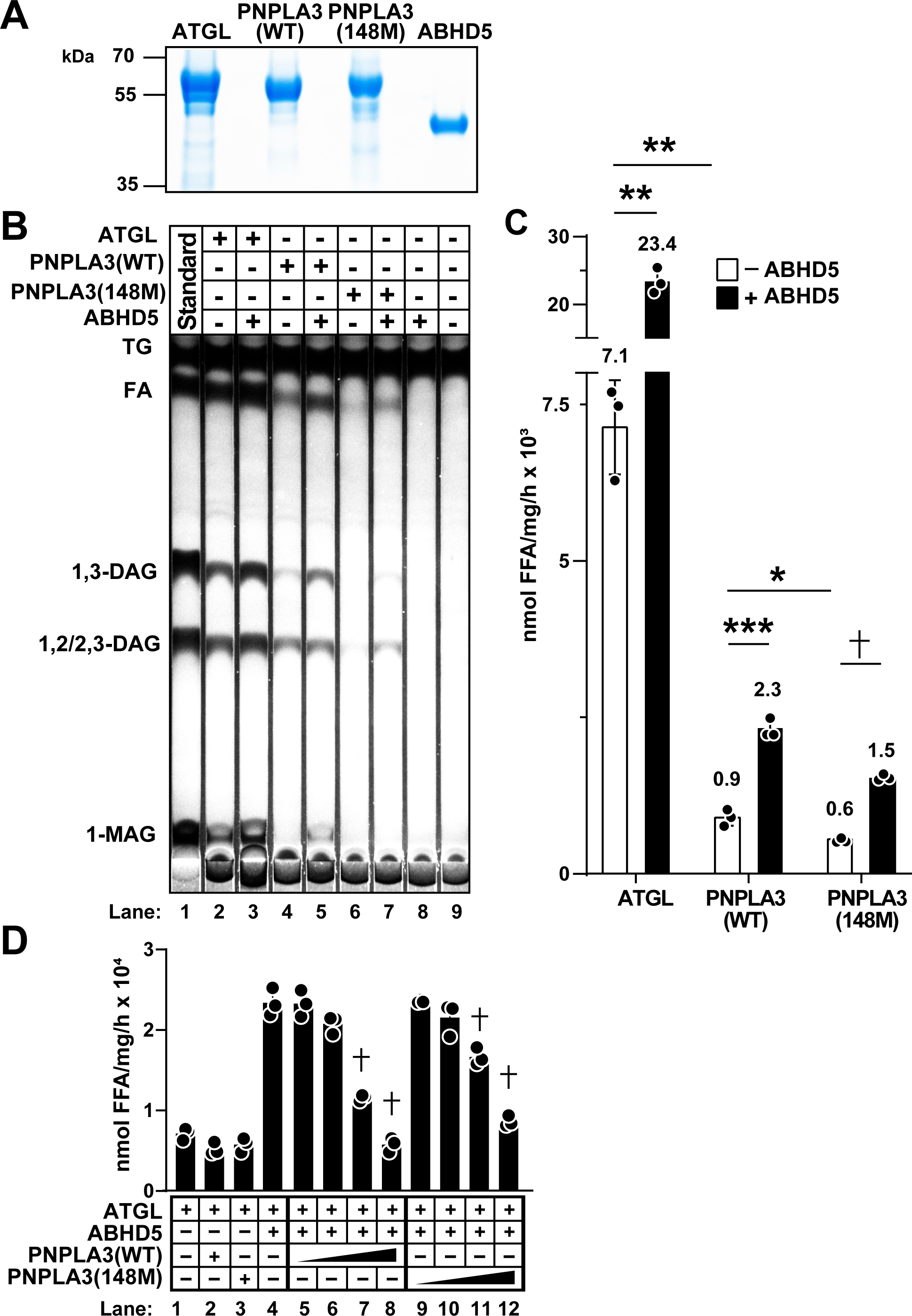
*In vitro* TG hydrolase activities of purified PNPLA3 and ATGL in the presence or absence of ABHD5. (A) Coomassie stain of recombinant human ATGL, PNPLA3(WT), PNPLA3(148M), and ABHD5 purified from mammalian HEK293S GnTI^-^ cells. ATGL, PNPLA3(WT) or PNPLA3(148M) (5 μg) and ABHD5 (3.5 μg) were subjected to SDS–PAGE. (B and C) Basal and ABHD5-activated TG hydrolysis of ATGL, PNPLA3(WT) and (148M) via (B) thin layer chromatography (TLC) and (C) gas chromatography-mass spectrometry (GC-MS). A total of 5 μg of purified ATGL, PNPLA3(WT), PNPLA3(148M) +/- an equimolar amount of ABHD5 were incubated for 1 h with lipid emulsions containing 6.68 mM triolein. Lipids were extracted and fractionated by TLC. Standards for the lipids were loaded in lane 1. (C) Free fatty acids (FA) were derivatized in tri-ethylamine (1%) and pentafluorobenzyl bromide (1%) in acetone and then quantified by GC-MS. Numbers above the bars represent values of means in each group. (D) Competition between PNPLA3 and ATGL for ABHD5 *in vitro*. Various molar ratios (from 0.5X to 3X relative to ATGL) of PNPLA3(WT) or (148M) were added to mixtures containing equimolar amount of ATGL and ABHD5. The protein mixtures were incubated with 25 μl of emulsion containing 6.68 mM triolein for 1 h at 37℃. The released FAs were quantified using GC-MS. Data represent mean ± SD; n=3 biological replicates; p values were determined by two-tailed t-tests with Welch’s correction or one-way ANOVA followed by Dunnett’s multiple comparisons test. *p<0.05, **p<0.01, ***p<0.001, †p<0.0001. This experiment was repeated twice and the results were similar.

To more precisely quantity and compare the TG hydrolase activities of ATGL and PNPLA3, we used gas chromatography-mass spectrometry (GC-MS) to measure rates of FA accumulation (Fig. 6C). The lipase activity of ATGL was 8-fold higher than that of PNPLA3. A ∼3.3-fold increase in free FA levels was seen with addition of ABHD5 to the reaction containing ATGL and a ∼2.5-fold increase was seen when ABHD5 was added to PNPLA3(WT) and PNPLA3(148M). We concluded from these experiments that PNPLA3, as well as ATGL, is activated by ABHD5 and that this effect occurs in the absence of other proteins.

To determine if PNPLA3 can compete with ATGL for ABHD5 activation *in vitro*, we incubated lipid emulsions with ATGL plus ABHD5 alone (1:1 molar ratio) or in the presence of either PNPLA3(WT) or PNPLA3(148M) and measured the effect on FA generation. A dose-dependent reduction in TG hydrolase activity was observed with addition of both forms of PNPLA3. From this experiment we concluded that the interaction between ABHD5 and ATGL is compromised in a similar fashion by the presence of PNPLA3(WT) and PNPLA3(148M). These results are similar to what we observed when we performed a competition assay in cultured cells (Fig. 1C).

## Conclusions

In this paper we provide evidence that PNPLA3(148M) alters TG mobilization in hepatocytes by sequestering ABHD5, the co-factor of ATGL. Using a variety of approaches, we tested directly the ABHD5 sequestration model. First, luciferase reconstitution assays were used in cultured hepatocytes to show that ABHD5 binds preferentially to PNPLA3 over ATGL (Fig. 1C) and that PNPLA3 competes with ATGL for binding to ABHD5 (Fig. 1E). Immunofluorescence was used to show that PNPLA3 was more effective than ATGL at recruiting/retaining ABHD5-LBD on LDs (Fig. 2C). In these experiments, we found no evidence that the 148M substitution altered the strength of binding of PNPLA3 to ABHD5. We documented previously that PNPLA3(148M) accumulates to much higher levels (40X) on hepatic LDs than PNPLA3(WT),^25^ and therefore we predicted that PNPLA3(148M) would bind and sequester more ABHD5. If the hepatic steatosis in *Pnpla3^M^*^/*M*^ mice is due to a paucity of ABHD5, then increasing levels of the co-activator would be predicted to rescue the steatotic phenotype. That was indeed what we observed when we overexpressed ABHD5 in *Pnpla3^M/M^* mice (Fig. 5D). These findings, when taken together, are consistent with the notion that PNPLA3(148M) causes steatosis by accumulating to high levels on LDs where it binds and sequesters ABHD5, thus compromising the activity of ATGL.

A possibility that we entertained prior to performing these studies was that PNPLA3(148M) promotes TG accumulation in the liver because it binds more strongly to ABHD5 than does PNPLA3(WT). However, we found no evidence to support this notion. PNPLA3(148M) and PNPLA3(WT) bound ABHD5 with similar characteristics when expressed together with ABHD5 in cells (Fig. 1C). Moreover, we found no differences in the ability of purified PNPLA3(WT) and PNPLA3(148M) to inhibit ABHD5-activated TG hydrolysis by ATGL (Fig. 6D). These results support the model that PNPLA3(148M)-induced steatosis is due to the high levels of PNPLA3(148M) on LDs rather than an increased affinity of the protein for ABHD5. Our results are largely consistent with those of Granneman and his colleagues^26^ who used luciferase reconstitution and co-immunoprecipitation to show that ABHD5 and PNPLA3 physically interact in cultured cells expressing both recombinant proteins, as well as in mouse liver and brown adipose tissue.^26^

To our knowledge, we have provided the first demonstration that ABHD5 directly activates the TG hydrolase activity of PNPLA3 (Fig. 6B), as it does ATGL.^18^ More detailed studies using purified components are ongoing to further characterize and compare the kinetics and substrate specificity of these two lipases and determine the relative effects of ABHD5 on these reactions.

We performed an extensive series of studies to map the interface(s) between PNPLA3 and ABHD5, reasoning that inhibition of the interaction would rescue the associated steatosis. We used AlphaFold to predict potential sites of physical association and these predictions were tested by making serial deletions and mutations in both proteins. All mutations that reduced interaction between the two proteins resulted in mislocalization or destabilization, presumably due to misfolding (data not shown). Other groups have implicated particular residues or regions of ABHD5 or ATGL as being essential for co-activation, but none have been confirmed to be necessary and sufficient.^22, 27–30^ The mechanism for ABHD5-mediated activation of ATGL remains to be determined. Elucidation of the atomic structure of the proteins holds the promise for shedding light on this important biological activity.

Johnson *et al*.^6^ reported that PNPLA3, like ATGL, is a TG hydrolase with specificity for TGs containing polyunsaturated fatty acids (PUFA). These results are consistent with our prior findings that *Pnpla3^-/-^*mice accumulate TG-PUFAs in hepatic LDs.^31^ Also consistent with our prior findings, they provide evidence that PNPLA3 mobilizes TG-PUFAs and enriches the PUFA content of phospholipids. They attribute the fatty liver phenotype of PNPLA3(148M) to a dearth of PUFAs in the phospholipids, resulting in reduced hepatic secretion of TG. In contrast to their results, we find that mice expressing PNPLA3(148M) have increased levels of phospholipid-PUFAs in hepatic LDs.^31^ Moreover, we (and others) have failed to observe any decrease in VLDL-TG secretion, in *Pnpla3^-/-^* mice^7, 8^ or any reduction in plasma TG levels in *Pnpla3^M/M^*mice.^9, 31^ We cannot rule out the possibility that the seemingly contradictory results of the two groups are due to differences in diet.

Johnson *et al.*^6^ also reported that both knockdown and knockout of PNPLA3 caused hepatic steatosis in mice. This finding has important mechanistic and clinical implications. If the phenotype of the *Pnpla3^-/-^* mice recapitulates that of the *Pnpla3^148M/M^*strain, it would be difficult not to conclude that the 148M substitution confers a loss-of-function mutation and that the steatosis associated with PNPLA3(148M) is a direct result of decreased catalytic activity. Arguing against this hypothesis is the finding that *Pnpla3^-/-^* mice have hepatic TG content levels that are similar to WT animals, even when challenged with a high-carbohydrate diet.^7, 8^ Moreover, inactivation of *Pnpla3* in a genetically obese (*ob/ob*) strain of mice failed to result in any increase in hepatic TG.^7^

Despite the PNPLA3(148M) variant causing steatosis through a gain-of-function mechanism, the effect requires that the TG hydrolase activity of PNPLA3 be reduced or ablated. Mice in which the catalytic serine is replaced with alanine (S47A), thus inactivating the enzyme, are a phenocopy of the *Pnpla3^M/M^* mice.^25^ However, a reduction or absence of PNPLA3 enzymatic function is not sufficient to cause hepatic steatosis. PNPLA3(148M) and PNPLA3(47A) must be expressed to observe the phenotype.^9, 25^ The mice only develop hepatic steatosis when subjected to diets that promote secretion of insulin (e.g., high-sucrose, high-fructose diets), which increases PNPLA3 expression.^10, 25^ These features are all compatible with the variants conferring a gain-of-function.

In a recent clinical trial in humans, knockdown of PNPLA3 using an anti-PNPLA3 siRNA led to a reduction in hepatic TG content in PNPLA3(148M) homozygotes.^12^ These findings are similar to what has been observed in mice; knockdown of PNPLA3(148M) mRNA using siRNAs or reduction of PNPLA3(148M) protein levels using proteolysis targeting chimeras both cause in a reduction in hepatic TG levels.^10, 11^ Experiments are now in progress to further define the molecular basis for the inhibition of ATGL activity by PNPLA3.

## Abbreviations

ABHD5: α/β hydrolase domain containing protein 5
AAV: adeno-associated virus
ATGL: adipose triglyceride lipase
CANX: calnexin
GC-MS: gas chromatography-mass spectrometry
HSD17B13: hydroxysteroid 17-β dehydrogenase 13
LBD: LD binding defective
LD: lipid droplet
L, LgBiT: LargeBiT
MDH: monodansylpentane
PNPLA3: patatin-like phospholipase domain-containing protein 3
PUFA: polyunsaturated fatty acids
S, SmBiT: SmallBiT
TG: triglyceride
TLC: thin layer chromatography
WT: wildtype.

## Data availability statement

All data presented in this manuscript will be made available upon request.

## Acknowledgements

We wish to thank Amy Whitaker, Paige Halverson, Panyun Wu, Christina Zhao, Fang Xu, Serena Banfi, Samantha Tomko and Tommy Hyatt for their excellent technical support. We thank Emma Bergman for her graphic contributions to the manuscript.

