## Supplementary Data for "PNPLA3(148M) promotes hepatic steatosis by interfering with triglyceride hydrolysis through a gain-of-function mechanism"

### Materials and Methods

*Materials*

Please see Table S1.

*Mice*

*Pnpla3* 148M knock-in mice (*Pnpla3$^{M/M}$* mice) were generated as described previously[1]. Ls-*Atgl$^{-/-}$* mice were generated by crossing *Atgl*-flox mice (B6N.129S-*Pnpla2$^{tm1Eek}$*/J, strain #:024278)), which have loxP sites flanking exon 2 through 7 of *Atgl*[2], and Albumin-Cre transgenic mice (strain 018961) provided by Philipp Scherer (UTSW). Animals were maintained on a 12 h light (7 am - 7 pm)/12 h dark (7 pm - 7 am) cycle and fed a chow diet (Teklad Mouse/Rat Diet 7001 or PicoLab Rodent Diet 5053) *ad libitum*[3]. For dietary challenge studies, mice were fed a high fructose diet (TD.89247, Teklad; 60% fructose) for 4 weeks. Mice were metabolically synchronized for 3 days with 18 h fasting and 6 h refeeding and sacrificed after the last feeding cycle unless indicated[1]. All animal experiments were performed with the approval of the Institutional Animal Care and Research Advisory Committee at the University of Texas Southwestern Medical Center in Dallas, Texas.

*Cell lines*

QBI-293A cells were growth in 6-well plates in complete medium [high sucrose DMEM plus 5% fetal calf serum (FCS)] in 8.8% $CO_2$ at 37ºC. For transient transfections, cells were treated with plasmids (2 µg) plus FuGENE 6 according to the manufacturer's instructions. Oleate (200 µM) was supplemented in  complete medium when needed. HuH7 cells were cultured in complete medium contained 10% FCS. GenJet In Vitro DNA Transfection Reagent was used to transfect HuH7 cells according to the instructions of the manufacturer. *ATGL* was inactivated in QBI-293A cells using CRISPR Cas-9 technology[4]. Briefly, guide RNAs targeting exon 2 of human ATGL gene were designed using the following website: https://www.benchling.com/crispr. The oligonucleotide guide pairs (see Table S1) were annealed and inserted into pX459 v2.0 prior to being used to transfect QBI-293A cells. Cells were then grown in complete medium supplemented with puromycin (0.5 μg/ml) for 48 h. Surviving cells were diluted to 0.5 cell/well in 96-well plates. After 2-3 weeks, single clones were isolated and tested for expression of ATGL by immunoblotting. Cells subcloned for three passages were maintained as ATGL knockout cell lines.

*Protein-protein interaction (NanoBiT) assay*

Complementary DNAs encoding the proteins of interest were inserted into NanoBiT system vectors (see Table S1). HuH7 cells were seeded in 96-well plates at a density of $1\times10^4$ cells/well. On day 1, cells were transfected with plasmids (0.2 μg) using 0.6 μl GenJet. After 21-24 h, the medium was replaced with 100 μl fresh medium and the cells were incubated for 1 h at 37ºC. A total of 25 μl/well of freshly made NanoGlo substrate (25 μl) was added to each well.  After 15 min at 37ºC, the plate was equilibrated to room temperature for 5 min before measuring the luminescence using an EnVision plate reader (Perkin Elmer). Cells were rinsed with PBS and cell lysis buffer (10 μl) plus protease inhibitor and benzonase was added to each well. Samples were pooled, mixed with Laemmli SDS-Sample Buffer, and then heated to 95ºC for 5 min before being subjected to SDS-PAGE (4-15%). Luminescent signals were measured and normalized to protein levels, as determined using LI-COR analysis of immunoblots.

*Immunoblotting and quantification*

Equal amounts of either LD proteins or cell lysates were subjected to SDS-PAGE. Proteins were transferred to nitrocellulose membranes and immunoblot analysis was performed using the indicated antibodies (see Table S1) and SuperSignal West Pico PLUS Chemiluminescent Substrate. Images of the immunoblots were scanned using an Odyssey FC Imager (LI-COR) and the expression levels of specific proteins were quantified using Image Studio Lite Ver 5.2 (LI-COR).

*Immunofluorescence microscopy*

Cells expressing genes of interest were fixed and the proteins were visualized using indicated antibodies as described[3]. Briefly, cells were rinsed twice with PBS and then fixed on slides using 4% paraformaldehyde for 15 min. The fixed cells were rinsed thrice with PBS and then incubated with 50 mM $NH_4Cl$ for 15 min to reduce autofluorescence. Cells were permeabilized with Triton X-100 (1%) for 2 min and then rinsed with PBS. Nonspecific epitopes were blocked by incubating the slides with fish skin gelatin (FSG) (0.2%) for 15 min and then with with indicated primary antibodies diluted in FSG (0.2%) overnight at 4ºC. Cells were then rinsed with PBS and incubated with secondary antibodies conjugated with fluorescent labels diluted in FSG (0.2%) for 35 min at room temperature. At The secondary antibody was then added at the same time that the LDs were stained using monodansylpentane (MDH) or LipidTOX Deep Red Neutral Lipid Stain (1:1,000 dilution) for 35 min. Cells were then rinsed with PBS and immersed into a drop of mounting medium on a slide pre-washed with ethanol and water. Samples were visualized using a confocal microscope (Zeiss LSM 880 or 800 or Leica TCS SP5). The images were analyzed using Fiji[5]. Briefly, different channels of images were split and merged. Cells were outlined using the freehand tool. The outlines were copied to other channels using Analyze > Tools > ROI Manager. To define the area of protein co-localized with LD, the threshold of LD and target proteins was set up. For LD localization of target protein, the threshold of LD channel was set up using Image > Adjust > Threshold and Otsu method was selected. The threshold of target protein was set up using the same method as LD except the cutoff was adjusted until top 10% values were selected. The LD and target protein channels were overlapped using Process > Image Calculator. The two channels were chosen and the operation "AND" was selected. The area of signals of the overlapped channel and target protein channel was measured separately (select the target cell, Analyze > Analyze Particles > set size to 0-Infinity and circularity to 0.00-1.00 > summarize > Total Area). The ratio of protein co-localized with LD was calculated by dividing the area of overlapped channel to the area of target protein channel. To quantify LD area, the threshold of LD channel was set up using Otsu method (Image > Adjust > Threshold). The cells were outlined and LD area in the target cell was measured in pixels (Freehand tool to select cell > Analyze > Measure > IntDen/255). The area was then converted to $\mu m^2$ by using area in pixels (IntDen/255) times pixel width times pixel height (pixel size: Image > Properties).

*Adeno-associated virus and adenovirus infection of mice*

Adeno-associated viruses (AAV8) with an albumin (1.9) promoter expressing mouse ABHD5 with three FLAG tags at the C-terminus or enhanced GFP were purchased from Vector Biolabs.

Mice were infected with AAVs [$1.5\times10^{11}$ (genomic copies) GC/mouse] in 200 µl saline by tail vein injection using a 1 ml syringe and 30G x ½ needle. Four weeks after infection, mice were metabolically synchronized for three cycles as described above and sacrificed after the last feeding cycle.

Adenoviruses expressing no insert (RR5), PNPLA3(WT) or PNPLA3(148M) with V5-His×6 tags at the C-terminus were constructed as described[6]. Mice were injected with $1.5\times10^{11}$ recombinant adenovirus particles in 200 µl normal saline by tail vein injection and metabolically synchronized for three days. Mice were sacrificed after the last feeding cycle.

*Lipid droplet isolation*

Hepatic LDs were isolated and purified as described[1, 3]. Briefly, livers were homogenized in ice-cold Buffer A (20 mM Tricine, 250 mM sucrose, pH 7.8) and the postnuclear supernatants (PNS) were collected after centrifugation (500*g* for 10 min). The PNS was overlaid with Buffer B (20 mM HEPES, 100 mM KCl, 2 mM $MgCl_2$, pH 7.4) and then centrifuged at 10,000*g* for 1 h. LDs were collected and washed with Buffer B three times. LD proteins were isolated by adding cold acetone and then subjecting the mixture to centrifugation at 15,000*g* for 15 min. The proteins were washed with acetone/ethyl ether (50/50, v/v) and air dried. The dried protein pellet was dissolved in PBS, 2% (v/v) SDS and 1 M urea. Protein concentrations were measured using the BCA assay.

*Quantitative Real-Time PCR*

Total RNA was extracted from liver samples using an RNA extraction kit (RNeasy Plus Universal Mini Kit,) and the concentration measured using a NanoDrop microvolume spectrophotometer (Thermo Fisher Scientific). The quality of RNA was assessed by agarose gel electrophoresis. Selected mRNA were reverse transcribed into cDNAs and the levels were quantified by real-time PCR as described[7]. Briefly, first-strand cDNAs were synthesized from 2 μg of total RNA using random hexamer primers and reverse transcription reagents (see Table S1). The real-time PCR reaction was performed in a final volume of 20 μl containing 20 ng of cDNA, 167 nM of primers, and 10 μl of SYBR Green PCR Master Mix (see Table S1). All reactions were performed in duplicate. Levels of mouse cyclophilin B mRNA were used as internal controls for normalization of transcript levels.

*Lipid measurements*

Lipids were extracted from frozen liver samples (~100 mg) using Folch solution and measured by enzymatic colorimetric assays[3, 8]. Briefly, tissues that had been flash frozen in liquid nitrogen and stored at -80ºC were homogenized using a Bead Ruptor Elite (OMNI International) in 1 ml Folch solution. The homogenates were transferred to glass tubes and Folch solution (4 ml) plus PBS (1 ml) was added. Tubes were vortexed for 30 s and centrifuged at 1,500*g* for 10 min. The lower phase was collected and Folch solution was added to a final volume of 5 ml. Samples (40 μl) were transferred to a 96-well polypropylene plate and 10 μl of chloroform-Triton mix (1:1) was added. The samples plus standards were dried at 37ºC, mixed with 300 μl of TG reagent (see Table S1) and then incubated at 37ºC for 20 min. The samples were transferred to a 96-well plate and the $OD_{515/735}$ ratios were measured. All values were normalized to the tissue weight.

*Protein expression and purification*

A cDNA encoding human PNPLA3 (NCBI Reference Sequence: NM_025225.3) and ATGL (NCBI Reference Sequence: NM_020376.4) were inserted into pEG BacMam with a FLAG tag added to the N-terminus. The coding sequences were validated by Sanger sequencing and then the plasmids were used to transfect mammalian HEK293S GnTI$^-$ cells[9]. After 8 h at 37°C, sodium butyrate was added to a final concentration of 10 mM and the temperature was reduced to 30°C. After 72 h, cells were homogenized in buffer D (20 mM HEPES, pH 7.5, 150 mM NaCl) supplemented with leupeptin (10 μg/ml) and PMSF (1 mM) and then lysed by sonication. Cell debris was removed by low-speed centrifugation and the supernatant was incubated with 1% (w/v) n-Dodecyl-β-D-Maltopyranoside (DDM) for 1 h at 4°C. The insoluble fraction was removed by centrifugation (15,000 rpm, 4°C, 30 min) and the supernatant was loaded onto an anti-FLAG M2 resin (Sigma) that had been pre-equilibrated with buffer D plus 0.02% DDM. The resin was washed sequentially with 15 ml of Buffer D containing the following additions: 0.02% DDM, 0.02% DDM plus 2 mM ATP and 4 mM $Mg^{2+}$, and 0.02% DDM by gravity flow. The lipase was eluted in 7.5 ml elution buffer (20 mM HEPES, pH 7.5, 150 mM NaCl, 0.02% DDM, and 0.1 mg/ml 3×FLAG peptide) and further purified by size-exclusion chromatography using a Superose 6 Increase, 10/300 GL column pre-equilibrated with buffer D plus 0.02 % DDM. Peak fractions containing the target protein were concentrated and stored in -80°C for further analysis.

For purification of human ABHD5 (NCBI Reference Sequence: NM_001355186.2), the same procedure was used except that an N-terminal Strep tag (WSHPQFEK) generated by PCR was added to the protein and the protein was purified using a Strep-Tactin XT 4Flow resin (IBA). The resin was washed without adding ATP and $Mg^{2+}$ and the recombinant ABHD5 was eluted from the column using biotin (50 mM).

*Lipolysis assay and TLC analysis*

Triolein was emulsified in phospholipids (PC/PI, 3:1, w/w) and FA free BSA (5% w/v) prepared in 0.1 M potassium phosphate buffer (KPB, pH 7.0) was added[10]. Purified PNPLA3, PNPLA3(148M) or ATGL (5 µg) were mixed with 25 µl of the emulsion (6.68 mM triolein) in a total volume of 50 µl for 1 h at 37°C. For ABHD5-stimulated TG hydrolase activity, the same amount of enzyme was mixed in a 1:1 molar ratio with ABHD5 (3.5 µg). As a negative control, ABHD5 and Buffer D plus 0.02% DDM were incubated with the emulsion. The reaction was terminated by adding 5 ml chloroform/methanol (2:1). After addition of 1 ml PBS, the mixture was vortexed for 30 s and centrifuged at 3,300 rpm for 5 min. The aqueous phase was discarded, and the solvent was evaporated under nitrogen. The dried lipids were dissolved in 100 µl chloroform/methanol (2:1) and fractionated by TLC using hexane/ethyl ether/acetic acid (120:30:8, v/v/v). The TLC plate was dried and stained with iodine, and then imaged using AlphaView SA (AlphaImager HP).

*GC-MS free fatty acid analysis*

Samples were prepared as described above for TLC analysis. The dried lipids were incubated with 50 µl of pentafluorobenzyl bromide (1%) and 50 µl of trimethylamine (1%) in acetone at room temperature for 1 h. Sample aliquots (25 µl) were transferred to a new GC vial and isooctane (975 µl) was added. A total of 1 µl of the mixture was subjected to GC/MS using an Agilent 7890/5975C GC-MS (Santa Clara, CA) by Negative Chemical Ionization equipped with an Rxi-5Sil-MS column (40 m × 0.180 mm; 0.18 µm film thickness). The hydrogen (carrier gas) flow rate was 2.0 ml/min, and the injection port temperature was 280°C. The initial oven temperature was 150°C and then the temperature was progressively increased by 15°C/min to a final temperature of 300°C. The temperature was maintained at 300°C for 3 min. The total run time was 15.0 min. The FAs were analyzed in selected ion-monitoring mode and the data was normalized to an internal standard (C16:1) prior to processing using Mass Hunter software (Agilent). All samples were injected and analyzed at least 3 times.

*Quantification and statistical analysis*

Differences among groups were analyzed using two-tailed t-test with Welch's correction or one-way ANOVA followed by Dunnett's or Sidak's multiple comparisons test or by 2-way ANOVA followed by Tukey's multiple comparisons test using GraphPad Prism 10 (GraphPad Software) or R Statistical Software (version 4). Data with non-normal distributions (e.g., luminescence values) were logarithmically transformed prior to performing ANOVA. Data were shown as mean ± SD unless otherwise stated. P-values less than 0.05 were considered statistically significant. The statistical details of experiments are provided in the Figure Legends.

**Supplemental Figures**

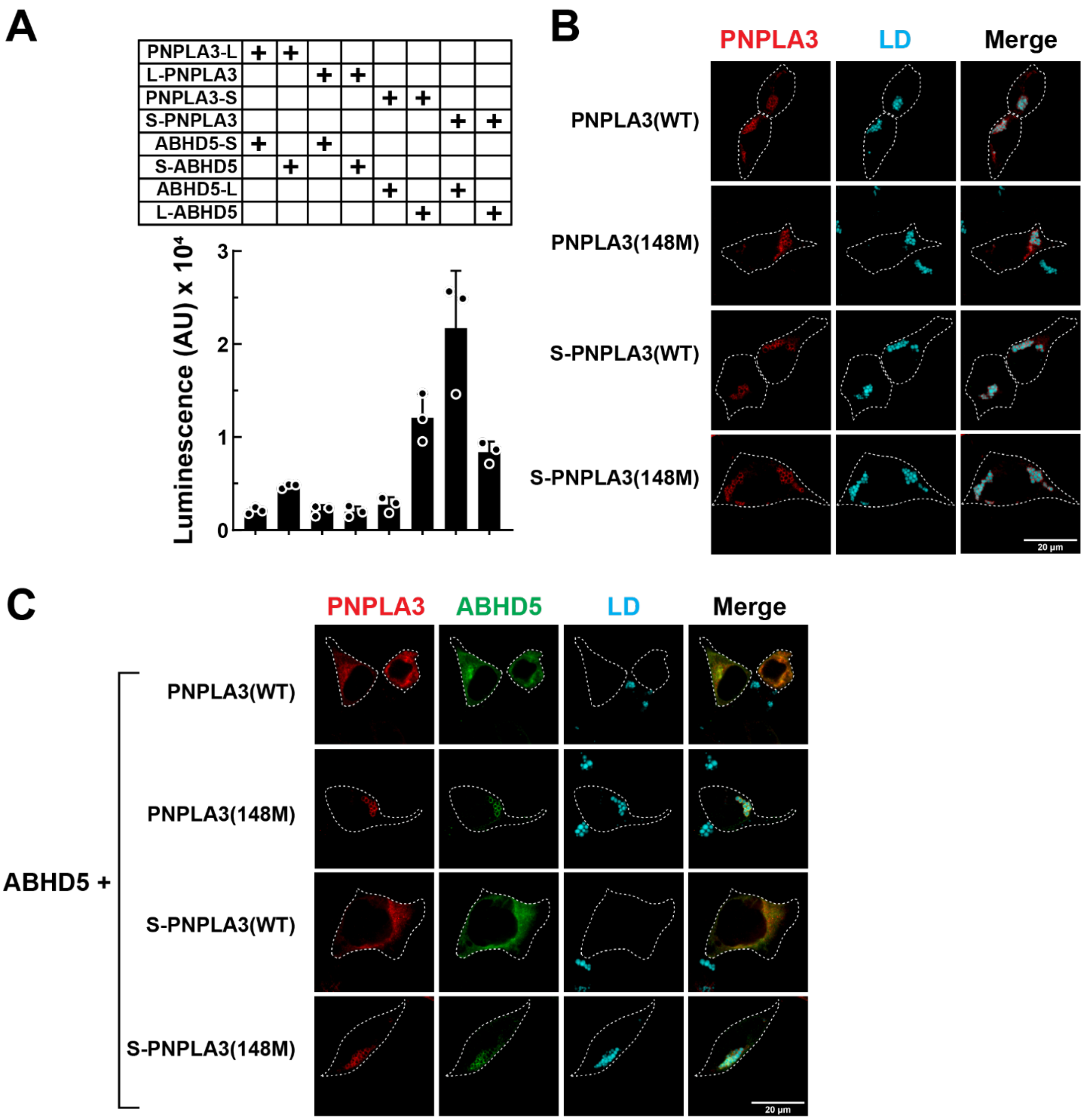

**Fig. S1. Effect of NanoBiT tag placement on expression and localization of PNPLA3 and ABHD5.** (A) LgBiT (L) and SmBiT (S) tags were placed at the N- or C-terminus of PNPLA3 and ABHD5. The proteins were expressed in a pairwise fashion in cells and the luminescence intensity was measured n=3 replicates/group. All proteins were expressed under TK promoter to avoid false positive caused by high protein expression levels. The experiment was repeated once, and the result was similar. AU: arbitrary unit. (B) Effect of adding SmBiT to the N-terminus of PNPLA3(WT) and PNPLA3(148M) on immunolocalization of the protein in QBI-293A cells grown in complete medium supplemented with oleate (200 μM). S-PNPLA3(WT) and S-PNPLA3(148M) were expressed under CMV promoter in a sufficient level for detection by immunofluorescence using anti-PNPLA3. LDs were stained using monodansylpentane (MDH). Scale bar: 20 μm. The addition of the SmBiT to the N-terminus of PNPLA3 did not alter its localization to LDs or activate its TG hydrolase activity. (C) Effect of addition of SmBiT to the N-terminus of PNPLA3(WT) and PNPLA3(148M) on cellular localization of the protein when it is co-expressed with ABHD5-myc. Immunofluorescence was performed using mouse anti-PNPLA3 monoclonal antibody and rabbit anti-myc polyclonal antibody as described in the Materials and Methods. LDs were stained using LipidTOX Deep Red (cyan). The patterns of expression of PNPLA3 and LDs were similar in cells receiving the tagged proteins and untagged proteins. Scale bar: 20 μm.

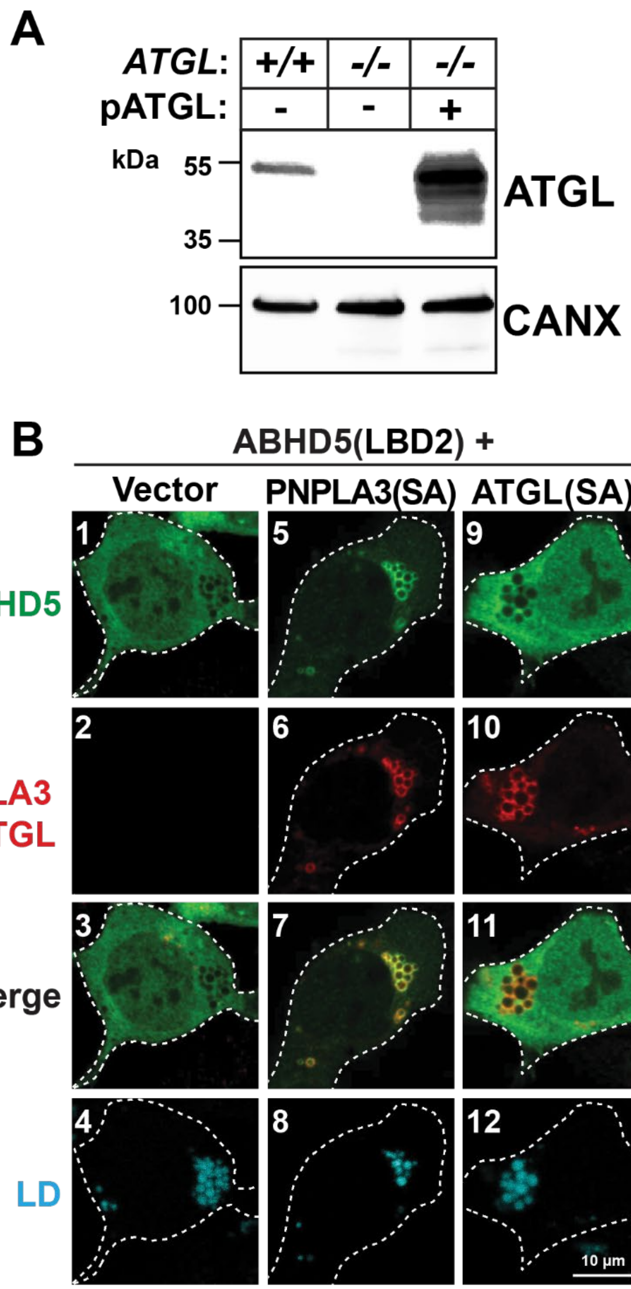

**Fig. S2. Comparison of the proportion of ABHD5-LBD2 that localizes to LDs containing PNPLA3 or ATGL in *ATGL*$^{-/-}$ QBI-293A cells.**

(A) Immunoblot result confirmed that *ATGL* had been successfully inactivated in QBI-293A cells using CRISPR-Cas9 as described in the *Materials and Methods*. *ATGL* knockout cells overexpressing ATGL(WT) served as a positive control. pATGL: plasmid ATGL. CANX, calnexin. (B) Recombinant ABHD5-LBD2-myc was co-expressed with empty vector,

PNPLA3(47A)-V5 or ATGL(47A)-V5 in *ATGL*$^{-/-}$ QBI-293A cells. Immunofluorescence was performed using mouse anti-V5 monoclonal antibody and rabbit anti-myc polyclonal antibodies as described in the Materials and Methods. LDs were stained using LipidTOX Deep Red. Green: ABHD5; Red: PNPLA3 or ATGL; Cyan: LD; Merge: ABHD5 signal merged with PNPLA3 or ATGL signal. White dashes outline the transfected cells. Scale bar: 10 µm.

**Supplementary Table**

**Table S1. Key Resources**

| REAGENT or RESOURCE | SOURCE | IDENTIFIER |
|---|---|---|
| Antibodies | | |
| Human PNPLA3 (11C5) | BasuRay et al.[11] | N/A |
| Mouse PNPLA3 (19A6) | BasuRay et al.[11] | N/A |
| ABHD5 | Novus | H00051099-M01 |
| ATGL | Cell Signaling Technology | 2138 |
| Anti-FLAG M2 Affinity Gel | Millipore-Sigma | A2220 |
| PLIN2 | Abcam | ab108323 |
| V5 | Thermo Fisher Scientific | R960-25 |
| myc | Abcam | ab9106 |
| Peroxidase AffiniPure Goat Anti-Rabbit IgG (H+L) | Jackson Immunoresearch Laboratories | 111-035-144 |
| Peroxidase AffiniPure Donkey Anti-Mouse IgG (H+L) | Jackson Immunoresearch Laboratories | 715-035-150 |
| Goat anti-Rabbit IgG (H+L) Secondary Antibody, Alexa Fluor™ 488 | Invitrogen | A-11034 |
| Goat anti-Mouse IgG (H+L) Secondary Antibody, Alexa Fluor™ 555 | Invitrogen | A-21422 |
| Bacterial and virus strains | | |
| Ad-RR5 | He *et al.*[6] | N/A |

| | | |
|---|---|---|
| Ad-PNPLA3(WT) | He *et al.*[6] | N/A |
| Ad-PNPLA3(148M) | He *et al.*[6] | N/A |
| AAV-ALB1.9-GFP | Vector Biolabs | SKU: VB1586 |
| AAV-ALB1.9-ABHD5[#] | Vector Biolabs | SKU: AAV-251823 |
| In-Fusion® Snap Assembly Master Mix with Competent Cells | Takara | 638952 |
| MAX Efficiency™ DH5α Competent Cells | Thermo Fisher Scientific | 18258012 |
| E.coli DH10Bac competent cells | Thermo Fisher Scientific | 10361012 |
| Chemicals, peptides, and recombinant proteins | | |
| 3×FLAG peptide | GeneScript | RP21087 |
| Benzonase Nuclease | Millipore Sigma | E1014-25KU |
| Biotin | Millipore-Sigma | B4501-5G |
| Bovine Serum Albumin | Millipore Sigma | 9048-46-8 |
| cOmplete Mini EDTA-free Protease Inhibitor Cocktail | Roche-Sigma Aldrich | 11836170001 |
| Cultrex® Poly-D-Lysine | R&D Systems | 3439-200-01 |
| Dimethyl Sulfoxide (DMSO) | Millipore Sigma | D2650 |
| DMEM High Glucose Medium | Corning | 10-013-CV |
| Fatty Acid Free Bovine Serum Albumin (BSA) | Millipore-Sigma | A7030 |
| Fetal Bovine Serum (FBS) | Millipore Sigma | F0926 |
| Fish Skin Gelatin | Millipore Sigma | G7765 |
| FLAG-ATGL | This paper | N/A |
| FLAG-PNPLA3(WT) | This paper | N/A |

| | | |
|---|---|---|
| FLAG-PNPLA3(148M) | This paper | N/A |
| Freestyle 293 expression medium | Gibco | 12338018 |
| FuGENE® HD Transfection Reagent | Promega | E2311 |
| GenJet™ In Vitro DNA Transfection Reagent for HuH-7 Cells (Ver. II) | SignaGen Laboratories | SL100489-HUH |
| HCS LipidTOX™ Deep Red Neutral Lipid Stain | Invitrogen | H34477 |
| HEPES | Millipore-Sigma | H3375 |
| LB broth | Fisher Scientific | BP1426-500 |
| Leupeptin | Vivitide | ILP-4041 |
| Monodansylpentane (MDH, AUTODOT™ Visualization Dye) | abcepta | SM1000a |
| Opti-MEM™ I Reduced Serum Medium | Thermo Fisher Scientific | 31985070 |
| Phenylmethylsulfonyl fluoride (PMSF) | Millipore-Sigma | 93482 |
| Phosphatidylcholine (PC) from egg yolk | Millipore-Sigma | P-3556 |
| Phosphatidylinositol (PI) from soybean | Millipore-Sigma | P-0639 |
| Puromycin Dihydrochloride | Millipore Sigma | P8833 |
| Sf-900 III SFM medium | Gibco | 12658027 |
| Sodium butyrate | Millipore-Sigma | 303410 |
| Sodium Oleate | Millipore Sigma | O7501 |
| Strep-ABHD5 | This paper | N/A |
| Strep-Tactin®XT 4Flow® resin | IBA | 2-5010-010 |
| Superose 6 Increrase 10/300 GL column | Cytiva | 29091596 |
| Tricine | Sigma | T0377 |
| Triolein | Millipore-Sigma | T-7140 |

| | | |
|---|---|---|
| VECTASHIELD® HardSet™ Antifade Mounting Medium | Vector Laboratories | H-1400 |
| Commercial assays | | |
| Analtech Preadsorbent Silica gel HL (TLC plate) | Miles Scientific | P43911 |
| Nano-Glo® Live Cell Assay System | Promega | N2011 |
| pcDNA™3.1/V5-His TOPO™ TA Expression Kit | Invitrogen | K480001 |
| Phusion® High-Fidelity DNA Polymerase | New England Biolabs | M0530 |
| Pierce BCA Protein Assay Kit | ThermoFisher | 23225 |
| *Power* SYBR Green PCR master Mix | Applied Biosystems | 4368708 |
| QuikChange II XL Site-Directed Mutagenesis Kit | Agilent Technologies | 200522 |
| Random hexamers | Thermo Fisher Scientific | N8080127 |
| RNeasy Plus Universal Mini Kit | Qiagen | 73404 |
| SuperSignal West Pico Chemiluminescent Substrate | ThermoFisher | 34580 |
| TaqMan reverse transcription reagents | Applied Biosystems | N8080234 |
| Trans-Blot Turbo RTA Mini 0.2 µm Nitrocellulose Transfer Kit | Bio-Rad | 1704270 |
| Triglycerides Reagent | Fisher Scientific | TR22421 |
| Verichem Laboratories MATRIX PLUS CHEM. KIT (Triglyceride standard) | Fisher Scientific | NC9592194 |
| Experimental models: cell lines | | |
| HuH7 | Nakabayashi et al.[12] | N/A |
| QBI-293A | Obiogene, formerly Quantum Biotechnologies, CA | N/A |

| | | |
|---|---|---|
| *ATGL*$^{-/-}$ QBI-293A | This paper | N/A |
| *Sf9* insect cell | ATCC | CRL-1711 |
| HEK293S GnTI$^{-}$ | ATCC | CRL-3022 |
| Experimental models: organisms/strains | | |
| Mouse: B6N.129S-*Pnpla2*$^{tm1Eek}$/J | The Jackson Laboratory | IMSR_JAX:024278 |
| Mouse: *Atgl* Liver Specific Knockout (Ls-*Atgl*$^{-/-}$) | This paper | N/A |
| Mouse: *Pnpla3* 148M knock-in (*Pnpla3*$^{M/M}$) | Smagris et al.[1] | N/A |
| Oligonucleotides | | |
| Primers: CRISPR-Cas9 knockout of *ATGL* in QBI-293A: PNPLA2 (ENSG00000177666) Assembly 1 FWD: CACCGCGCCGACGTAGTAGACGCCG | This paper | N/A |
| Primers: CRISPR-Cas9 knockout of *ATGL* in QBI-293A: PNPLA2 (ENSG00000177666) Assembly 1 REV: aaacCGGCGTCTACTACGTCGGCGC | This paper | N/A |
| Primers: CRISPR-Cas9 knockout of *ATGL* in QBI-293A: PNPLA2 (ENSG00000177666) Assembly 2 FWD: CACCGACCCCGGTGACCAGCGCCG | This paper | N/A |
| Primers: CRISPR-Cas9 knockout of *ATGL* in QBI-293A: PNPLA2 (ENSG00000177666) Assembly 2 REV: aaacCGGCGCTGGTCACCGGGGTC | This paper | N/A |
| qPCR primers: mouse Cyclophilin B: TGGAGAGCACCAAGACAGACA | This paper | N/A |
| qPCR primers: mouse Cyclophilin B: TGCCGGAGTCGACAATGAT | This paper | N/A |
| qPCR primers: mouse PNPLA3: CGAGGCGAGCGGTACGT | This paper | N/A |
| qPCR primers: mouse PNPLA3: TGACACCGTGATGGTGGTTT | This paper | N/A |

| | | |
|---|---|---|
| qPCR primers: mouse ABHD5: AATGTGTCCCCTGCACTTACAA | This paper | N/A |
| qPCR primers: mouse ABHD5: GAACATCAGCGTCCATATTCTGTT | This paper | N/A |
| qPCR primers: mouse ATGL: GAGAGAACGTCATCATATCCCACTT | This paper | N/A |
| qPCR primers: mouse ATGL: CCACAGTACACCGGGATAAATGT | This paper | N/A |
| qPCR primers: mouse PLIN2: GGTGATGGCAGGCGACAT | This paper | N/A |
| qPCR primers: mouse PLIN2: CGCCATCGGACACTTCCTTA | This paper | N/A |
| qPCR primers: human PNPLA3: TGCAACTTGCTACCCATTAGGA | This paper | N/A |
| qPCR primers: human PNPLA3: TCGCAATGGCAGATTCCA | This paper | N/A |
| Recombinant DNA | | |
| Plasmid: SmBiT-PNPLA3(WT/148M)-V5 | This paper | N/A |
| Plasmid: SmBiT-ATGL-V5 | This paper | N/A |
| Plasmid: SmBiT-HSD17B13-V5 | This paper | N/A |
| Plasmid: SmBiT-PRKACA | Promega | N2015- N204A |
| Plasmid: LgBiT-PNPLA3 | This paper | N/A |
| Plasmid: PNPLA3-LgBiT | This paper | N/A |
| Plasmid: PNPLA3-SmBiT | This paper | N/A |

| | | |
|---|---|---|
| Plasmid: LgBiT-ABHD5 | This paper | N/A |
| Plasmid: ABHD5-SmBiT | This paper | N/A |
| Plasmid: SmBiT-ABHD5 | This paper | N/A |
| Plasmid: pcDNA-SmBiT-PNPLA3(WT/148M) | This paper | N/A |
| Plasmid: HSD17B13-3×HA-3×FLAG-V5 | This paper | N/A |
| Plasmid: ATGL-3×HA -3×FLAG -V5 | This paper | N/A |
| Plasmid: PNPLA3(WT/148M) -3×HA -3×FLAG -V5 | This paper | N/A |
| Plasmid: pSpCas9(BB)-2A-Puro (PX459) V2.0 | Addgene (gift from Feng Zhang)[4] | 62988 |
| Plasmid: PNPLA3(WT, 148M or 47A)-V5 | Wang et al.[3] | N/A |
| Plasmid: ABHD5-WT/LBD1/LBD2-myc | This paper | N/A |
| Plasmid: PNPLA3(WT/148M)-LBD-V5 | This paper | N/A |
| Plasmid: ATGL | This paper | N/A |
| Plasmid: ATGL(47A)-V5 | Wang et al.[3] | N/A |
| Plasmid: pEG BacMam | Addgene (gift from Eric Gouaux)[9] | 160451 |
| Plasmid: pEG BacMam-FLAG-PNPLA3(WT/148M) | This paper | N/A |
| Plasmid: pEG BacMam-FLAG-ATGL | This paper | N/A |
| Plasmid: pEG BacMam-Strep-ABHD5 | This paper | N/A |
| Software and algorithms | | |
| Image Studio Lite v5.2 | LI-COR | N/A |
| R Statistical Software v4 | R Core Team | https://www.r-project.org/ |
| Prism 10 | GraphPad | N/A |

| | | |
|---|---|---|
| ImageJ (Fiji) | Schneider et al.[5] | https://imagej.net/ij/ |
| Other | | |
| Mouse diet: chow diet 7001 | Envigo | Teklad 7001 |
| Mouse diet: chow diet 5053 | LabDiet | 5053 |
| Mouse diet: high sucrose diet | MP Biomedicals | 901682 |
| Mouse diet: high fructose diet | Envigo | Teklad TD.89247 |
| EnVision Multilabel Plate Readers | Perkin Elmer | 2104-0010 |
| GC-MS | Agilent | 7890/5975C |
| Leica confocal microscope | Leica | TCS SP5 |
| Zeiss confocal microscope | Zeiss | LSM 800 or 880 |
